# LUBAC is required for RIG-I sensing of RNA viruses

**DOI:** 10.1101/2022.02.24.481814

**Authors:** Helena C Teague, Charlotte Lefevre, Eva Rieser, Diego de Miguel, Daniel Patricio, Marisa Oliveira, Daniel S Mansur, Nerea Irigoyen, Henning Walczak, Brian J Ferguson

## Abstract

The ability of cells to mount an interferon response to virus infections depends on intracellular nucleic acid sensing pattern recognition receptors (PRRs). RIG-I is an intracellular PRR that binds short double stranded viral RNAs to trigger MAVS-dependent signalling. The RIG-I/MAVS signalling complex requires the coordinated activity of multiple kinases and E3 ubiquitin ligases to activate the transcription factors that drive type I and type III interferon production from infected cells. The linear ubiquitin chain assembly complex (LUBAC) regulates the activity of multiple receptor signalling pathways in both ligase-dependent and -independent ways. Here, we show that the three proteins that constitute LUBAC have separate functions in regulating RIG-I signalling. Both HOIP, the E3 ligase capable of generating M1-ubiquitin chains, and LUBAC accessory protein HOIL-1 are required for viral RNA sensing by RIG-I. The third LUBAC component, SHARPIN, is not required for RIG-I signalling. These data cement the role of LUBAC as a positive regulator of RIG-I signalling and as an important component of antiviral innate immune responses.

## Introduction

The interferon (IFN) response is a potent method of restricting virus replication at the site of infection. The ability of cells in infected tissues to rapidly detect invading viruses and mount an IFN response is a critical determinant of the outcome of viral disease^1^. Many RNA viruses are sensed by the intracellular pattern recognition receptor (PRR) retinoic-acid induced gene-I (RIG-I), which shows binding preference for short, double stranded hairpin RNAs with a 5’ triphosphate group^2,3^. Upon ligand binding, RIG-I oligomerises via its caspase recruitment domains (CARDs) and binds the adaptor mitochondrial antiviral-signalling protein (MAVS)^4^. The oligomerisation of RIG-I and MAVS at the mitochondrial membrane leads to the formation of a large multi-protein signalling complex that co-ordinates RIG-I signalling outcomes^5^. The key output of RIG-I signalling is the transcription of IFN-I/III, cytokines and chemokines driven by the activation of the transcription factors nuclear factor-kappa B (NF-κB) and interferon regulatory factor 3 (IRF3) at the RIG-I/MAVS signalling complex (SC) ^6^.

Regulated assembly of the RIG-I/MAVS SC requires the coordinated activity of E3 ubiquitin ligases and kinases that ultimately recruit and activate NF-κB and IRF3^5,7,8^. IRF3 is activated by phosphorylation by the kinases TBK1 and IKKε, allowing phospho-IRF3 dimerisation and translocation to the nucleus^9^. NF-κB is activated by IKK complex-dependent phosphorylation and subsequent degradation of the inhibitor protein IκBα. This step releases active NF-κB allowing it to translocate to the nucleus. Active, nuclear NF-κB and IRF3 co-ordinate a IFN-I/III and inflammatory transcriptional signature by binding to the promoters of specific genes, either independently or in tandem^10^. During this process, activation and modification of multiple and likely redundant TRAF proteins, K63-chain generating E3 ligases, results in the recruitment of the IKK complex via the ubiquitin binding-domain of NEMO (IKKγ)^7,11^. TBK1 and IKKε are subsequently recruited, possibly independently and by mechanisms that involve binding to ubiquitinated proteins in the complex, resulting in IRF3 phosphorylation. The IKK complex phosphorylates IκBα and recruits the K48-chain E3 ligase β-TRCP to modify p IκBα, tagging it for degradation.

The linear ubiquitin chain assembly complex (LUBAC) is responsible for attaching M1 ubiquitin chains to target proteins^12^. LUBAC consists of three proteins, HOIL-1-interacting protein (HOIP), which encodes the M1-chain E3 ligase activity; Heme-oxidised IRP2 Ubiquitin ligase-1 (HOIL-1); and Shank-associated RH domain-interacting protein (SHARPIN)^13^. M1 ubiquitin chains regulate multiple immune signalling pathways^13^ and LUBAC is recruited to a number of receptor signalling complexes, including the TNFR1-SC^14^ and TLR3-SC^15^. LUBAC’s function in RIG-I signalling is unclear, with studies indicating that it positively, negatively and redundantly regulates RIG-I signalling outputs^7,16–20^. In this study we set out to identify the contribution of the individual LUBAC components and the E3 ligase activity of LUBAC to the antiviral innate immune response downstream of RIG-I. We establish here specific roles for the individual LUBAC components in RIG-I signalling and show that whilst HOIP and HOIL-1 are essential for RNA-virus driven interferon responses, SHARPIN is not.

## Materials and Methods

### Cells

All cell lines were grown in Dulbecco’s Modified Eagle’s Medium (DMEM) with 4.5 g/L D-glucose, 8mM L-glutamine and Sodium Pyruvate (Gibco), supplemented with 10% foetal calf serum (FCS, Pan Biotech) and 100 U/mL Penicillin and Streptomycin (Gibco). Cells were incubated at 37°C, 5% CO2 and 3% O2 in a humidified incubator.

### CRISPR/Cas9 editing

The human genomic sequences of DDX58 (encoding RIG-I) and SHARPIN were identified on ENSEMBL (http://www.ensembl.org/index.html: DDX58 (RIG-I) ENSG00000107201 and SHARPIN ENSG00000179526). Small guide (sg)RNAs were designed using Benchling (www.benchling.com). sgRNA sequences were DDX58: AAAGTCCAGAATAACCTGCA and SHARPIN: CCTAGTCCGAGGTGCCACCG. Guides were synthesised as forward and reverse complementary DNA oligonucleotides (IDT) with BbsI restriction sites, to enable annealing into the pSpCas9(BB)-2A-GFP (PX458) plasmid (Addgene #48138). A549 cells were transfected with PX458-containing plasmids and single-cells sorted by GFP positivity to generate clonal knockout (KO) lines. Successful KO’s were verified by immunoblotting.

### Viruses

SeV Cantell strain and Zika virus ZIKV/H.Sapiens/Brazil/PE243/2015 (ZIKV PE243) were used for infection experiments. To quantify ZIKV by plaque assay, samples were 10-fold serial diluted in serum-free DMEM. 400 μL of dilutions were added to Vero cells in duplicate and incubated for 1 hour at 37°C. The inoculum was removed and replaced with a 50:50 mix of 3% LMP agarose and 2x MEM 4% FCS. Cells were incubated for 5 days before being fixed overnight at room temperature using formal saline (4% formaldehyde, 0.9% sodium chloride, 90% H2O) and then stained with Toluidine Blue and plaques counted.

### RT-qPCR

RNA was extracted from treated cells with 250 μL lysis buffer and purified by spin column or phenol/chloroform extraction. 500 ug purified RNA was used for cDNA synthesis by Superscript III Reverse Transcriptase (Invitrogen). cDNA was diluted 1:3 in nuclease-free water (Ambion) and added to 5 μL SyGreen HiROX mix (PCR Biosystems) in 384 well plates with 0.5 μM forward and reverse primers (sequences in Supplementary table 1) run on a Viia7 Real-Time PCR machine (Thermo Scientific). Fold induction of the target gene was calculated relative to *GAPDH* in human cells and *Hprt* in murine cells.

### Immunoblotting

Whole cell lysates were prepared by lysing in approximately 100 μL per 2.5×10^6^ cells. The lysis buffer used for A549 was radioimmunoprecipitation assay (RIPA) buffer (50 mM Tris-HCl pH 8, 1% Nonidet P-40 (NP-40), 0.5% sodium deoxycholate, 0.1% SDS, 150 mM NaCl) and for murine embryonic fibroblasts (MEFs) was 30 mM Tris-HCl pH 7.4, 1% Triton X-100, 20 mM NaCl, 2 mM KCl, 2 mM EDTA and 10% glycerol with protease inhibitors (Roche) and phosphatase inhibitors (Sigma) where appropriate. Protein concentration was determined by bicinchoninic acid (BCA) assay (Thermo Scientific) to enable equal loading of protein samples. Gels were run in a Mini-PROTEAN system (BioRad), transferred onto nitrocellulose membrane and analysed with specific primary and secondary antibodies listed in Supplementary Table 2.

### ELISA

A DuoSetELISA assay (R&D) was used to detect the presence of human CXCL10/IP-10 in the supernatants of infected or stimulated A549 cells using TMB (Abcam) as the substrate solution and 0.3 M H_2_SO_4_ as the stop solution.

### Flow cytometry

Cells were washed twice in PBS, detached with trypsin and resuspended in DMEM 2.5% FCS. Cells were pelleted by centrifugation at 600 x g for 6 minutes and fixed in 100 μL per 1×10^6^ cells of PhosFlow Lyse/Fix buffer (BD Bioscience) at 37 °C for 10 minutes. Fixation was stopped by addition of 1 mL of PBS 1% FCS and cells were stored at 4 °C overnight in PBS. Cells were pelleted by centrifugation and further fixed and permeabilised in 1 mL of 88% methanol PBS at 4 °C for 30 minutes. Cells were washed 3 times in PBS 1% FCS, each time pelleted by centrifugation at 850 x g for 6 minutes at 4 °C. Cells were incubated with primary antibody diluted in PBS 1% FCS (25 μL per 1×10^6^ cells), for 1 hour at room temperature in the dark. Details of antibodies are in Supplementary Table 3. The washing steps were repeated, and cells were resuspended in filtered PBS in FACS tubes and stored at 4°C until analysis. Control samples, no antibody, single antibody and positive control samples, were also stained under the same conditions. Samples were analysed by flow cytometry using an Attune NxT Acoustic Focusing Cytometer (Fisher Scientific) and analysed in FlowJo Version 10.

### Viability assay

To quantify cell viability, a Nucleocounter NC-250 Vitality assay was used. Cells were washed twice with PBS, trypsinised and resuspended in DMEM 2.5% FCS to a total volume of 1 mL per 5×105 cells (1 well). The cell suspension was mixed with NC-250 Solution 6, containing VB-48 vitality dye and propidium iodide (PI), at a 20:1 dilution, and this was added to a NC-slide. Slides were loaded into the Nucleocounter NC-250 and PI / VB-48 fluorescence intensity was quantified to analyse cell viability.

### Co-Immunoprecipitation

A549 cells stably expressing tandem affinity purification (TAP)-tagged HOIP or NEMO re-introduced into their respective KO lines, were seeded in 15 cm dishes. Cells were stimulated with 100 U/mL IFNα and incubated for 24 hours before being infected with SeV for the indicated time. Cells were washed twice in 5 mL cold PBS and scraped in 0.5 mL lysis buffer 1 (100 mM NaCl, 40 mM Tris-HCl, pH 7.5, 1 mM CaCl2, 1 mM MgCl2) with protease inhibitors (Roche) and phosphatase inhibitors (Sigma). Cells were lysed for 30 minutes on ice and 40 minutes on a rotating wheel at 4 °C. Insoluble debris was pelleted by centrifugation at 16200 x g for 10 minutes at 4 °C and cleared lysate was transferred to a new Eppendorf tube. Pellets were resuspended in 0.5 mL lysis buffer 2 (100 mM NaCl, 40 mM Tris-HCl, pH 7.5, 1 mM CaCl2, 1 mM MgCl2, 1% Triton X-100 and 0.1% SDS) with protease inhibitors (Roche) and phosphatase inhibitors (Sigma) and twice subjected to sonication at 20 Hz for 10 seconds, with a minute on ice between. Sonicated samples were subjected to centrifugation at 16200 x g for 20 minutes at 4 °C. Cleared lysates from pre- and post-sonification were combined. 35 μL of cleared lysate was taken for an input sample and 7 μL of 6x loading buffer was added to the remainder. 25 μL per sample of Flag-M2 beads (Sigma), pre-washed once with PBS and 3 times with the respective lysis buffer, were added to the cleared lysates and this was incubated on a rotating wheel at 4 °C for 16 hours. Beads were pelleted by centrifugation at 2,400 x g for 5 minutes at 4 °C and unbound material was removed using a 1 mL needle with a 20-gage syringe. Beads were washed by addition of 1 mL of respective lysis buffer (without protease or phosphatase inhibitors) followed by centrifugation as before. The washing procedure was repeated four more times. 40 μL of 2x loading buffer with 330 mM DTT was added to beads. Immunoprecipitation samples were heated at 95 °C for 5 minutes and were analysed by Western blotting.

### Immunofluorescence staining

A549 (WT and HOIP-/-) cells were seeded onto a 13 mm cover slips in 24-wells plates overnight, followed by ZIKV infection for 24 hours or 100 ng/mL TNF stimulation for 30 minutes. Cells were then fixed in 3% paraformaldehyde for 20 minutes and permeabilizated with 0.1% Triton X-100 diluted in blocking buffer (2 % bovine serum albumin in PBS) for 5 minutes. Primary and secondary antibodies (listed in Supplementary table 4) were diluted in blocking buffer. Cells were incubated with primary and then secondary antibodies for 1 hour. Samples were washed in PBS between each step. Images were acquired on an Olympus BX41 microscope.

## Results

### RIG-I is the dominant RNA sensor in A549 cells

To help define the outputs of RIG-I activation and its signalling mechanisms, we first generated a RIG-I knockout A549 cell line using CRISPR/Cas9 editing (Fig. 1A). Infection of wild type (WT) A549 cells with Sendai virus (SeV) or Zika virus (ZIKV) resulted in robust IFN-I (*IFNB1*) and IFN-III (*IFNL1*) transcription and activation of both IRF3-dependent (*ISG54*)^21^ and NF-κB-dependent (*NFKBIA*)^22^ genes (Fig. 1). In SeV- and ZIKV-infected RIG-I KO cells there was an almost complete loss of IFN-I and IFN-III response and a failure to transcribe other IRF3 and NF-κB dependent genes (Fig. 1 B, C). The response to the RIG-I specific ligand 3p-hpRNA was also lost in RIG-I KO cells (Fig. 1D), although the IFN-response to this ligand was weak, possibly due to low transfection efficiency. The transcriptional response to transfection of the dsRNA mimetic poly(I:C) was also abrogated in RIG-I KO cells (Fig. 1E). Since poly(I:C) can be sensed intracellularly by both RIG-I and MDA5 as well as by TLR3 in endosomes, the loss of poly(I:C)-driven transcription in RIG-I KO cells suggested that little or no MDA5 or TLR3 activity is present in A549.

**Figure 1:**
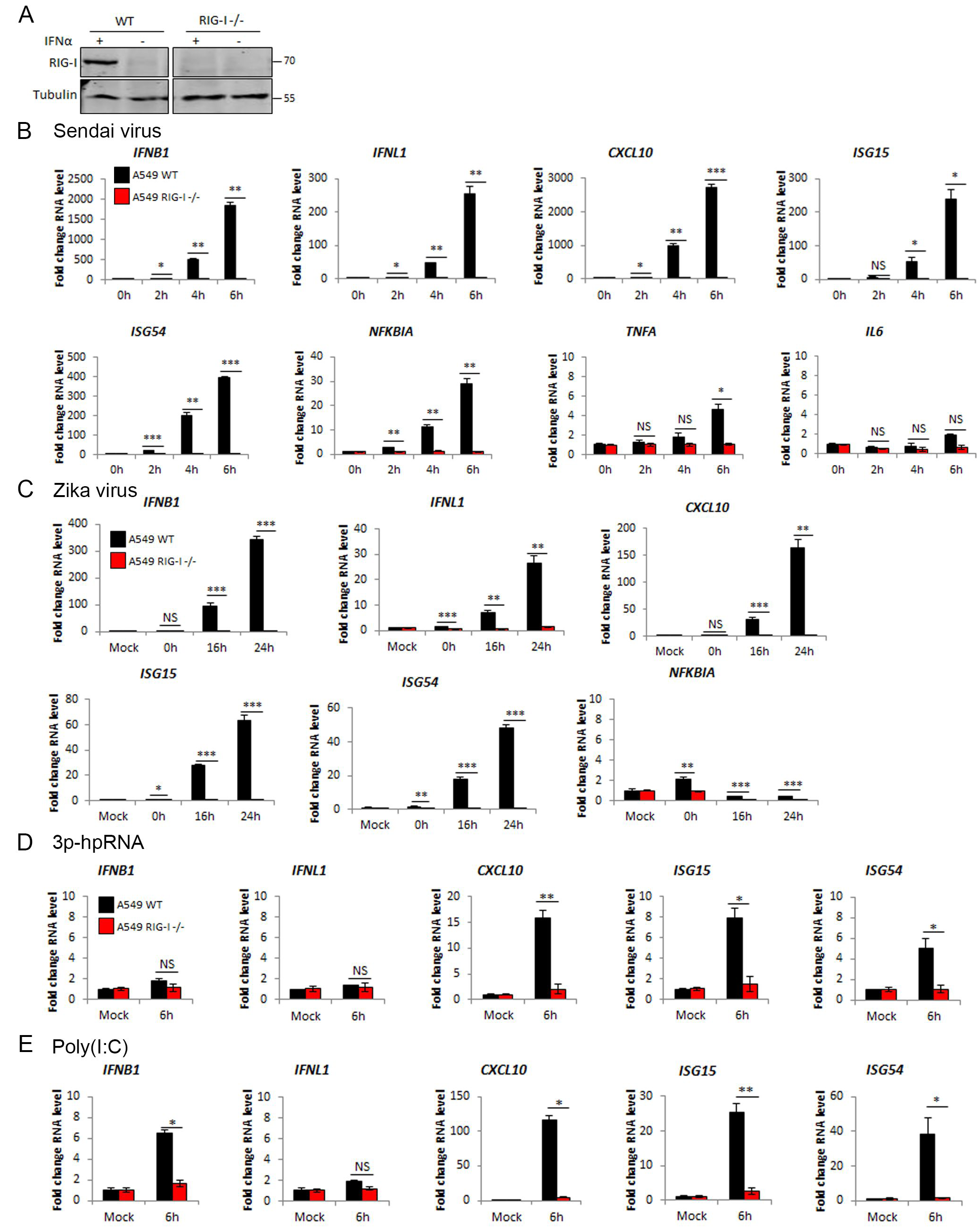
RIG-I dependent RNA and RNA virus sensing in A549 cells. A) Western blotting analysis of A549 WT and RIG-I -/- cells with and without stimulation with IFNα for 24 hours. qPCR to measure transcription of indicated genes in A549 WT and RIG-I -/- cells stimulated by B) SeV infection at 1:300 dilution, C) Zika virus infection at MOI 3, D) transfection with 1 μg 3p-hpRNA and E) transfection with 1 μg Poly(I:C).

### HOIP is required for anti-viral RIG-I signalling and for the IFN response to RNA viruses

Since the transcriptional response to SeV, ZIKV and intracellular synthetic RNAs was found to be dependent on RIG-I, we used this system to define the contribution of the E3 ligase HOIP to RIG-I signalling. In CRISPR/Cas9-generated HOIP KO A549 cells infected with SeV, transcription of *IFNB* and *IFNL* was >95% reduced compared to infected WT cells, indicating that HOIP is essential for the IFN-I and IFN-III responses to RNA virus infection (Fig. 2A). The loss of RIG-I dependent gene activation in HOIP KO cells extended to significant reductions in *CXCL10* and *ISG15*, as well as *ISG54* and *NFKBIA* transcription, indicating that both IRF3 and NF-κB-dependent responses to SeV were significantly impaired by loss of HOIP (Fig. 2A). Similar loss of transcription was observed in response to RNA transfection in HOIP KO cells (Fig. 2B). Analysis of intracellular signalling events showed that the activation of IRF3 and NF-κB signalling triggered by SeV infection in WT A549 cells was impaired in HOIP KO cells (Fig. 2C). TBK1, IRF3 and IκBα phosphorylation were all reduced at 2 and 4 h post infection, confirming that HOIP is required for the complete activation of both IRF3 and NF-κB pathways downstream of RIG-I signalling. HOIP KO cells were also defective in virus- and RNA-driven CXCL10 secretion (Fig. 2D), indicating loss of HOIP results in the overall loss of RIG-I signalling.

**Figure 2:**
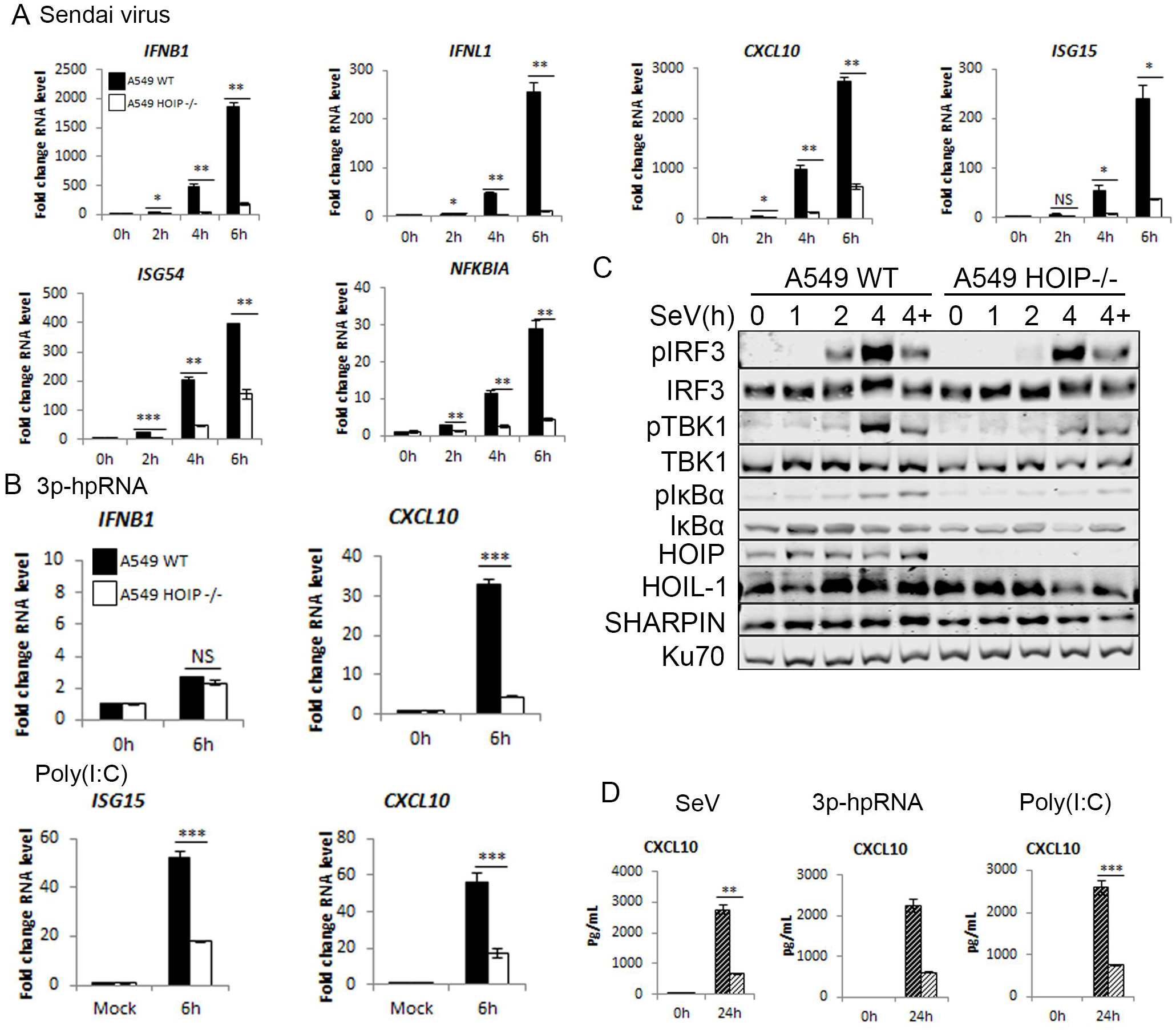
HOIP is required for RIG-I driven transcription, chemokine secretion and signalling pathway activation. A549 WT and HOIP -/- infected with SeV at 1:300 dilution and A) qPCR to measure transcription of indicated genes and B) Western blotting analysis in the presence and absence of 10 μM MG-132. C) qPCR to measure transcription of indicated genes in A549 WT and HOIP -/- cells transfected with 1 μg 3p-hpRNA or Poly(I:C). D) ELISA to measure CXCL10 secretion in A549 WT and HOIP -/- cells infected with SeV at 1:300 dilution or transfected with 1 μg 3p-hpRNA or 1 μg Poly(I:C).

As well as IFN responses, RIG-I signalling can result in regulated cell death^23,24^, therefore we quantified apoptotic and non-apoptotic death following SeV infection in WT and HOIP KO cells. Using phosFlow, we confirmed a reduction in phospho-IRF3 levels in HOIP KO cells compared to WT, but only observed active caspase-3 in 2% of cells during infection (Supplementary Fig. S1A). The lack of RIG-I driven caspase activation was corroborated by quantifying cell viability after infection. Cells infected with SeV showed ∼10% cell death in comparison with ∼70% cell death following staurosporine treatment, with no significant difference between WT and HOIP KO cells (Supplementary Fig. S1B). As such, in A549 cells RIG-I was not found to activate a regulated cell death pathway, irrespective of the presence or absence of HOIP.

To assess the impact of HOIP on infection with a replicating RNA virus, we infected WT and HOIP KO cells with ZIKV. *IFNB* and *IFNL* transcription was almost abrogated in HOIP KO cells compared to WT and there was an 80% reduction in *CXCL10* transcription (Fig. 3A). This defective transcriptional response was not a result of loss of infectivity or replicative capacity of ZIKV in HOIP KO cells as the virus titres and E protein expression levels were not impacted by loss of HOIP (Fig. 3B, C). We also assessed the impact of HOIP loss on IRF3 and NF-κB P65 nuclear translocation during ZIKV infection. In WT infected cells P65 and IRF3 were found translocated to the nucleus in multiple ZIKV infected cells. This translocation was lost in HOIP KO cells, consistent with the loss of cytoplasmic RIG-I signalling (Fig. 3D, E). HOIP is therefore essential for the transcriptional outputs of RIG-I signalling and for the IRF3 and NF-κB dependent IFN-I/III response to RNA virus infections.

**Figure 3:**
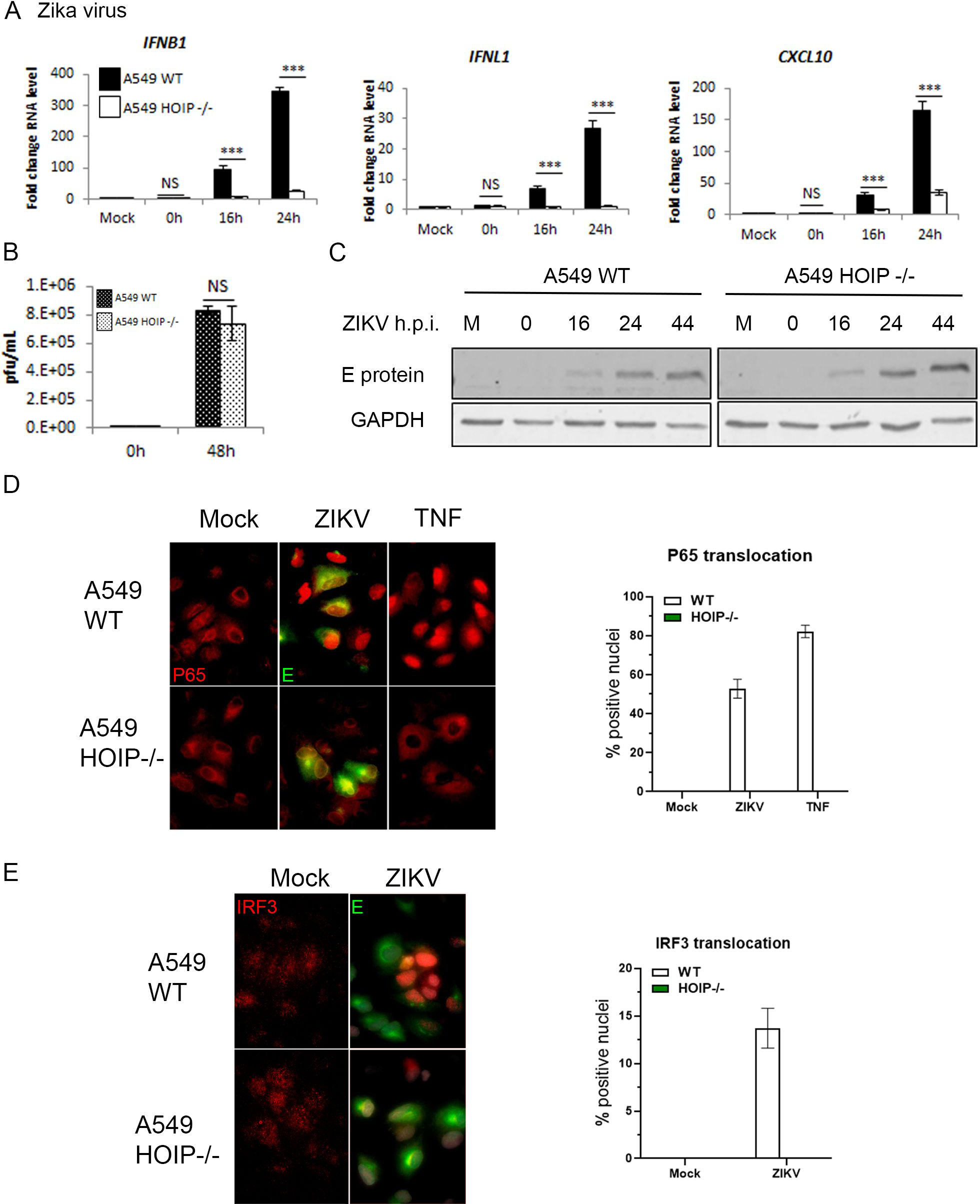
HOIP is required for ZIKV-driven interferon responses. A549 WT and HOIP -/- cells infected with ZIKV at MOI 3 and A) qPCR to measure transcription of indicated genes, B) ZIKV replication measured by plaque assay on Vero cells and C) Western blotting analysis. Quantification of nuclear translocation of D) NF-κB P65 and E) IRF3 in A549 WT and HOIP -/- cells infected with ZIKV at MOI 1 for 24 hours or stimulated with 100 ng/mL TNF, analysed by immunofluorescence (left panels) and quantified by scoring cells with nuclear staining (right panels).

### HOIL-1 is required for anti-viral RIG-I signalling

To understand the function of HOIL-1 in RIG-I signalling we initially attempted to create HOIL-1 KO A549 cells but failed to isolate knockout clones. We instead used MEFs, completely deficient in HOIL-1 expression (Supplementary Fig. S2). As complete HOIL-1 KO is embryonically lethal in mice, but can be partially rescued by backcrossing to TNF KO^25^, we used *Tnf*^*-/-*^*/Rbck1* ^*+/-*^ and *Tnf*^-/-^/*Rbck1*^-/-^ MEFs infected with SeV or transfected with synthetic RNAs. MEFs lacking HOIL-1 were found to be defective in RIG-I-driven IFN-I transcription and activation of both IRF3 and NF-κB signalling after SeV infection (Fig. 4A, B). Following RNA transfection, *Ifnb, Cxcl10, Isg54, Isg15, Nfkbia* and *Il6* transcription were also significantly reduced in HOIL-1 KO MEFs (Fig. 4C, D). This data defines HOIL-1, along with HOIP, as a key component of RIG-I signalling and indicates that LUBAC’s positive regulation of RIG-I signalling is conserved between human and murine cells.

**Figure 4:**
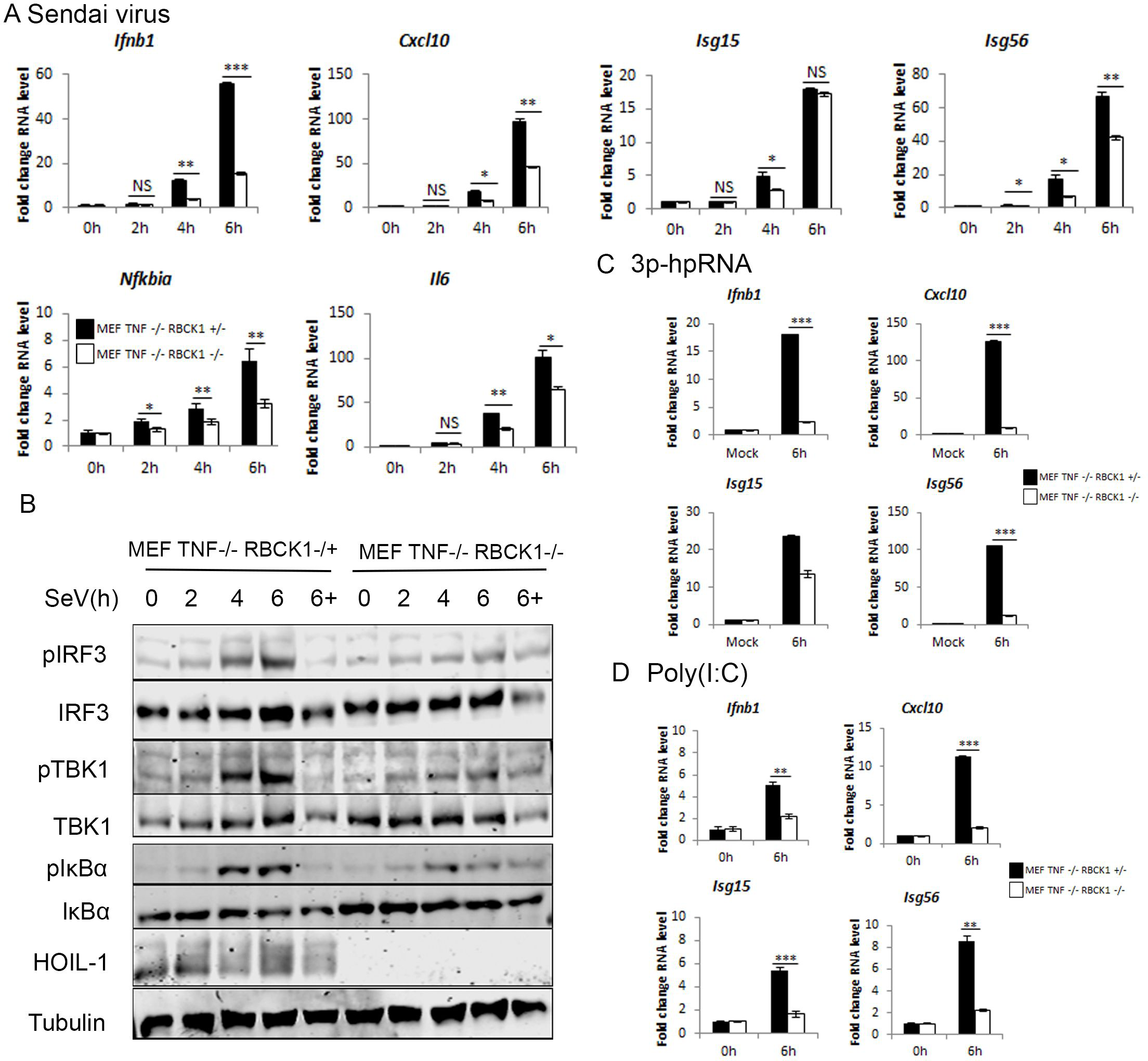
HOIL-1 is required for RIG-I immune response to SeV and synthetic RNAs. MEF TNF -/- Rbck +/- and TNF -/- Rbck -/- cells infected with SeV at a 1:300 dilution and A) qPCR to measure transcription of indicated genes and B) Western blotting analysis of signalling protein activation in the presence and absence of 10 μM MG-132. qPCR to measure transcription of indicated genes in MEF TNF -/- HOIL +/- and TNF -/- HOIL -/- cells transfected with C) 1 μg 3p-hpRNA and D) 1 μg Poly(I:C).

### SHARPIN is not required for anti-viral RIG-I signalling

The third LUBAC component, SHARPIN, has no ligase activity and acts as a structural protein, co-ordinating LUBAC and its interactions with other protein complexes^26^. To analyse the impact of SHARPIN on RIG-I signalling, we used two systems. We generated a SHARPIN KO A549 cell line (Fig. 5), and also used *cpdm* MEFs, which contain a germline mutation in the murine *Sharpin* gene that results in complete loss of SHARPIN protein expression^26^. Infection of SHARPIN KO A549 cells resulted in increased RIG-I-driven gene activation compared to WT cells, with interferon transcription as well as IRF3 and NF-κB-dependent gene transcription all increased in KO cells (Fig. 5A). In response to RNA transfection there was no significant alteration in *CXCL10* transcription, but a significant reduction in *ISG15* transcription (Fig. 5B) Analysis of the intracellular signalling events indicated that loss of SHARPIN did not impact TBK1, IRF3 or IκBα phosphorylation following SeV infection (Fig. 5C), but the increase in RIG-I signalling output is also observed at the level of CXCL10 protein secretion (Fig. 5D). Infection of *cpdm* MEFs, lacking SHARPIN expression (Supplementary Fig. S3A), with SeV resulted in a slight reduction in NF-κB-dependent genes, but increased IRF3-dependent transcription (Supplementary Fig. S3B). Transfection with RIG-I-specific RNA ligand, however, showed no impact of SHARPIN expression on RIG-I-driven transcription (Supplementary Fig. S3C). Poly(I:C)-driven transcription was impaired in *cpdm* MEFs (Supplementary Fig. S3D), although this might be explained by interference from active TLR3 or MDA5 signalling in MEFs^15^. Overall there was clear evidence that SHARPIN is not required for RIG-I signalling in humans or mice, and evidence that SHARPIN loss results in increased IRF3-dependent transcription under certain conditions. Comparison of the relative contribution of the three separate LUBAC components therefore defines both HOIP and HOIL-1 as being essential for RIG-I signalling and for the IFN-response to virus infections, but SHARPIN as dispensable for this process.

**Figure 5:**
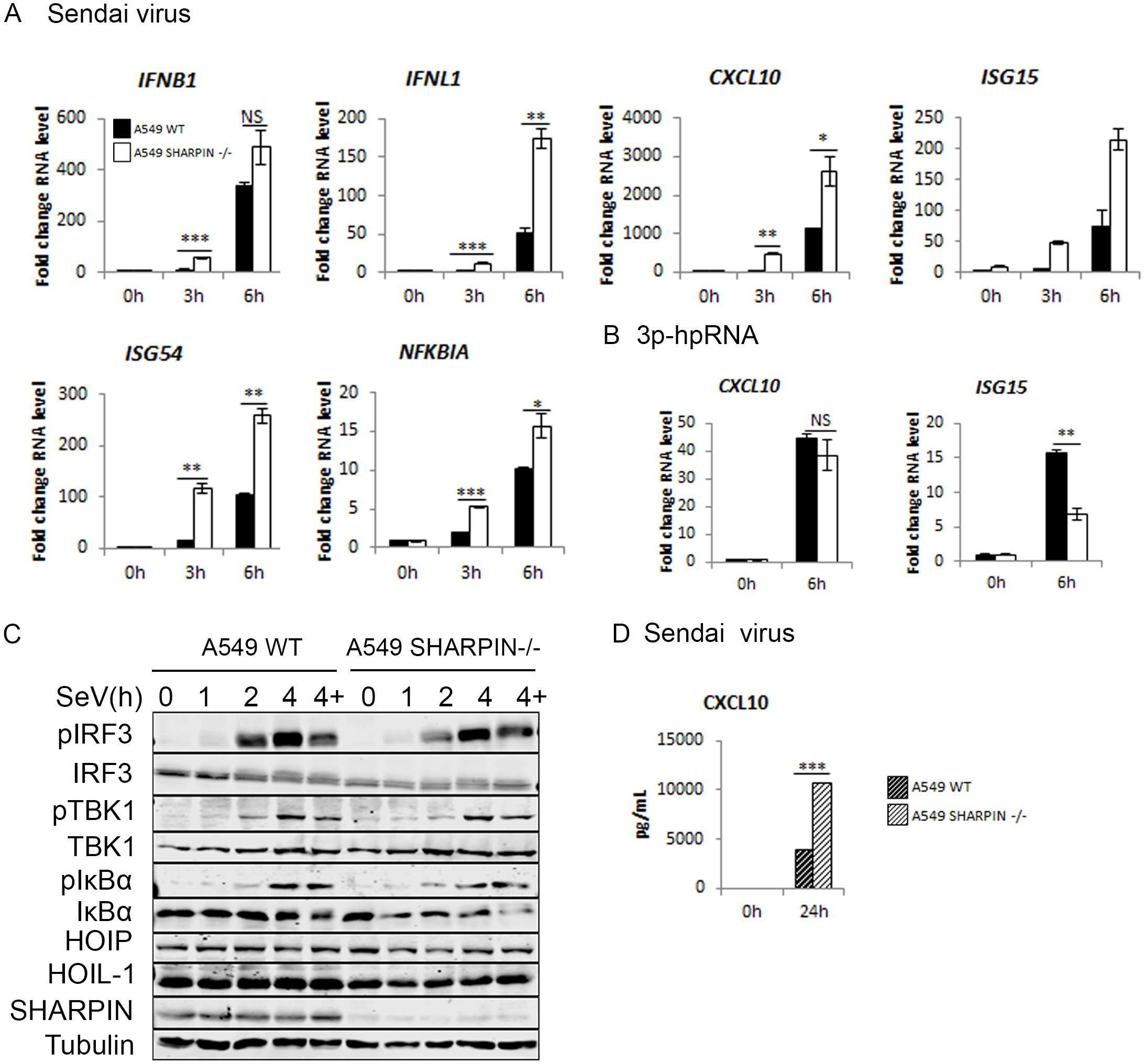
SHARPIN is not required for RIG-I immune response to SeV and synthetic RNAs in A549 cells. A549 WT and SHARPIN -/- cells infected with SeV at a 1:300 dilution and A) qPCR to measure transcription of indicated genes, B) Western blotting analysis of signalling protein activation in the presence and absence of 10 μM MG-132 and C) CXCL10 secretion measured by ELISA. qPCR to measure transcription of indicated genes in A549 WT and SHARPIN cells transfected with D) 1 μg 3p-hpRNA and E) 1 μg Poly(I:C).

### HOIP E3 ligase activity is partially required for RIG-I dependent IFN production

To further understand the function of LUBAC in RIG-I signalling, we asked whether the E3 ligase activity of HOIP contributes to RIG-I signalling. We used HOIP KO cells with a tandem-affinity purification (TAP)-tagged HOIP or the single point mutant TAP-HOIP-C885S lacking E3 ligase activity^27^. TAP-HOIP and TAP-HOIP-C885S were expressed at similar amounts but at a higher level than endogenous HOIP in WT A549 cells (Fig. 6A). SeV infection of WT, HOIP KO, TAP-HOIP-WT and TAP-HOIP-C885S rescue showed that TAP-HOIP-WT fully rescued the IFN-I/III response to SeV infection and inactivation of HOIP’s E3 ligase activity resulted in significantly less *IFNB, IFNL* and *CXCL10* transcription and CXCL10 secretion, when compared with WT A549 cells or HOIP KO cells rescued with WT TAP-HOIP (Fig. 6B, D). There was no observable difference in SeV-driven TBK1, IRF3 or IκBα activation between cells expressing TAP-HOIP or TAP-HOIP-C885S (Fig. 6C), indicating that HOIP’s E3 ligase activity is not required for RIG-I signalling activation. In response to synthetic RNA transfection, TAP-HOIP rescued cells transcribed significantly more *IFNB* and *CXCL10* than TAP-HOIP-C885S cells, confirming the phenotype shown during SeV infection (Fig. 6E). This data indicates that HOIP has a dual role in regulation of RIG-I signalling, being both dependent and independent of the E3 ligase function.

**Figure 6:**
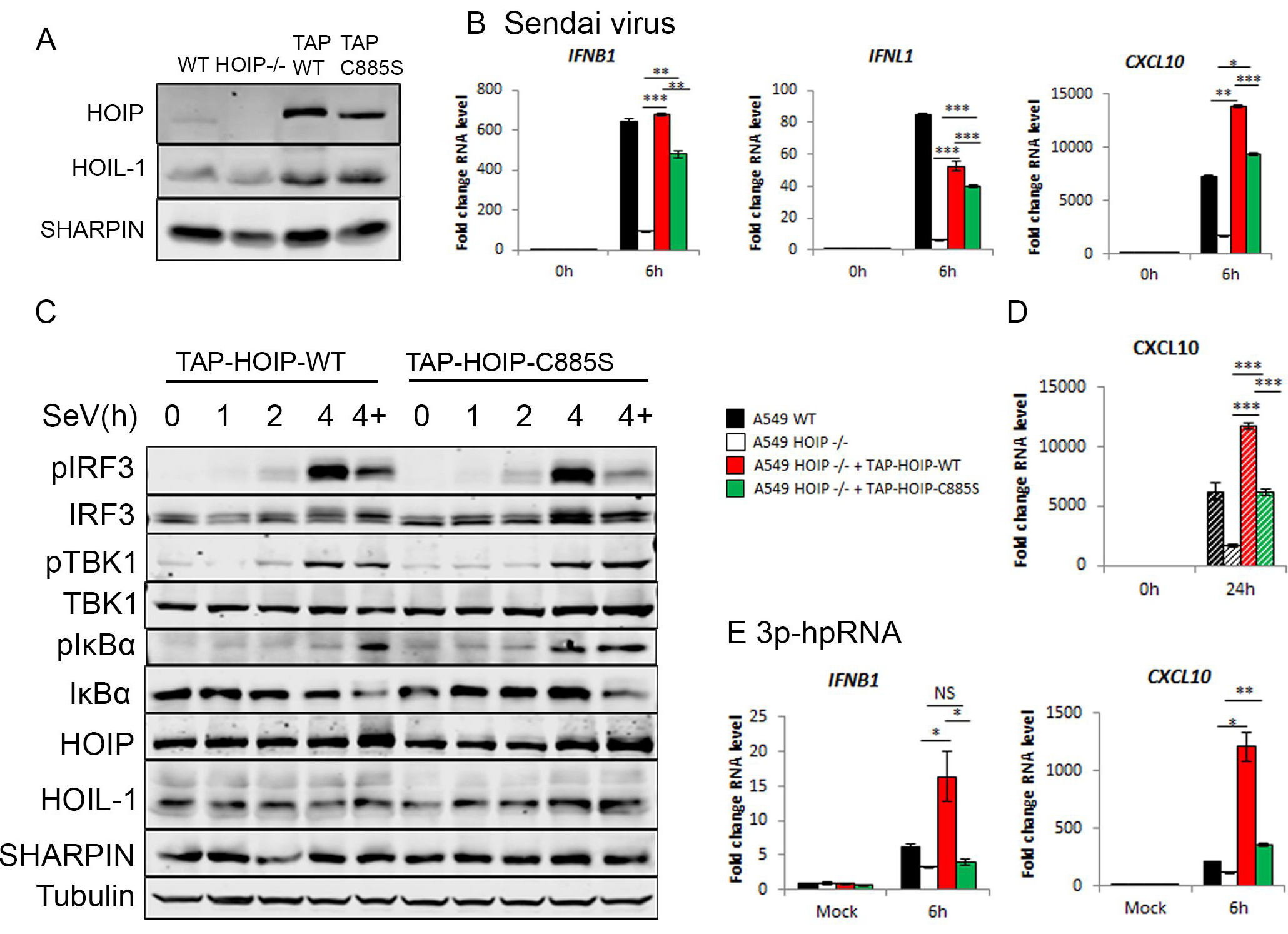
The E3 ligase activity of LUBAC is partially required for its function in RIG-I signalling. A) Western blotting analysis of A549 WT, HOIP -/-, TAP-HOIP-WT and TAP-HOIP- C885S cells. A549 WT, HOIP -/-, TAP-HOIP-WT and TAP-HOIP-C885S cells infected with SeV at a 1:300 dilution and B) qPCR to measure transcription of indicated genes and C) ELISA to measure CXCL10 secretion. D) Western blotting analysis of signalling protein activation in A549 TAP-HOIP-WT and TAP-HOIP-C885S cells infected with SeV at a 1:300 dilution in the presence and absence of 10 μM MG-132. qPCR to measure transcription of indicated genes in A549 WT, HOIP -/-, TAP-HOIP-WT and TAP-HOIP-C885S cells transfected with E) 1 μg 3p- hpRNA and F) 1 μg Poly(I:C).

### NEMO and TBK1/IKKε are essential for IFN-I responses to RIG-I stimulation

To further probe the mechanisms downstream of RIG-I that may co-ordinate with LUBAC to activate IFN responses and to assess tools for analysing RIG-I signalling complexes we analysed further A549 KO lines. As expected, MAVS KO cells were unable to mount a transcriptional response to SeV infection, confirming that MAVS is essential for RIG-I signalling in A549 cells^6^ (Supplementary Fig S4A, B). Using NEMO KO cells^28^, we also found that NEMO is essential for the IFN-response triggered by RIG-I signalling^29^, as NEMO KO cells infected with SeV or transfected with RNAs failed to transcribe *IFNB, IFNL*, or *CXCL10*, but this response could be fully rescued by re-expression of TAP-NEMO in the KO cells (Fig. 7A, Supplementary Fig. S4C,D). In NEMO KO cells we observed residual *NFKBIA* transcription, indicating that NEMO contributes to, but is non-essential for RIG-I-driven NF-κB activation (Fig. 7A). Downstream of PRRs, the kinases TBK1 and IKKε are necessary for IRF3 signalling^9^. We analysed the potential redundancy of these kinases and their contribution to IRF3 and NF-κB activation following RIG-I stimulation. Individual KO of TBK1 or IKKε^28^ had only minor impacts on gene activation following SeV infection or synthetic RNA transfection (Fig 7B and Supplementary Fig. S4E-G). Knockout of both TBK1 and IKKε, however, resulted in abrogation of IFN- and IRF3-dependent gene transcription, although had no impact on *NFKBIA* transcription, indicating that TBK1 and IKKε contribute redundantly to IRF3 activation and are not required for NF-κB activity downstream of RIG-I signalling (Fig. 7B). To confirm these observations, we analysed activation of TBK1, IRF3 and IκBα activation in NEMO, TBK1, IKKε and TBK1/ IKKε KO A549 cells (Fig. 7C). In NEMO KO cells, IRF3 and TBK1 phosphorylation were nearly abrogated following SeV infection and, although IκBα phosphorylation was maintained, the protein was not degraded in NEMO KO cells (Fig. 7C), consistent with the partial defect in NF-κB-dependent transcription (Fig. 7A). NEMO therefore functions as an essential regulator of RIG-I-driven IRF3 activation and is partially required for IκBα activity. Similarly, IRF3 phosphorylation following SeV infection was maintained in TBK1 and IKKε single KO cell lines, but in the TBK1/IKKε KO cells, IRF3 phosphorylation was abrogated (Fig. 7D) whilst IκBα phosphorylation and degradation was unaffected (Fig. 7D). Therefore, TBK1 and IKKε act redundantly downstream of RIG-I to phosphorylate IRF3 and activate IFN-I/III transcription but are not required for RIG-I driven NF-κB activation.

**Figure 7:**
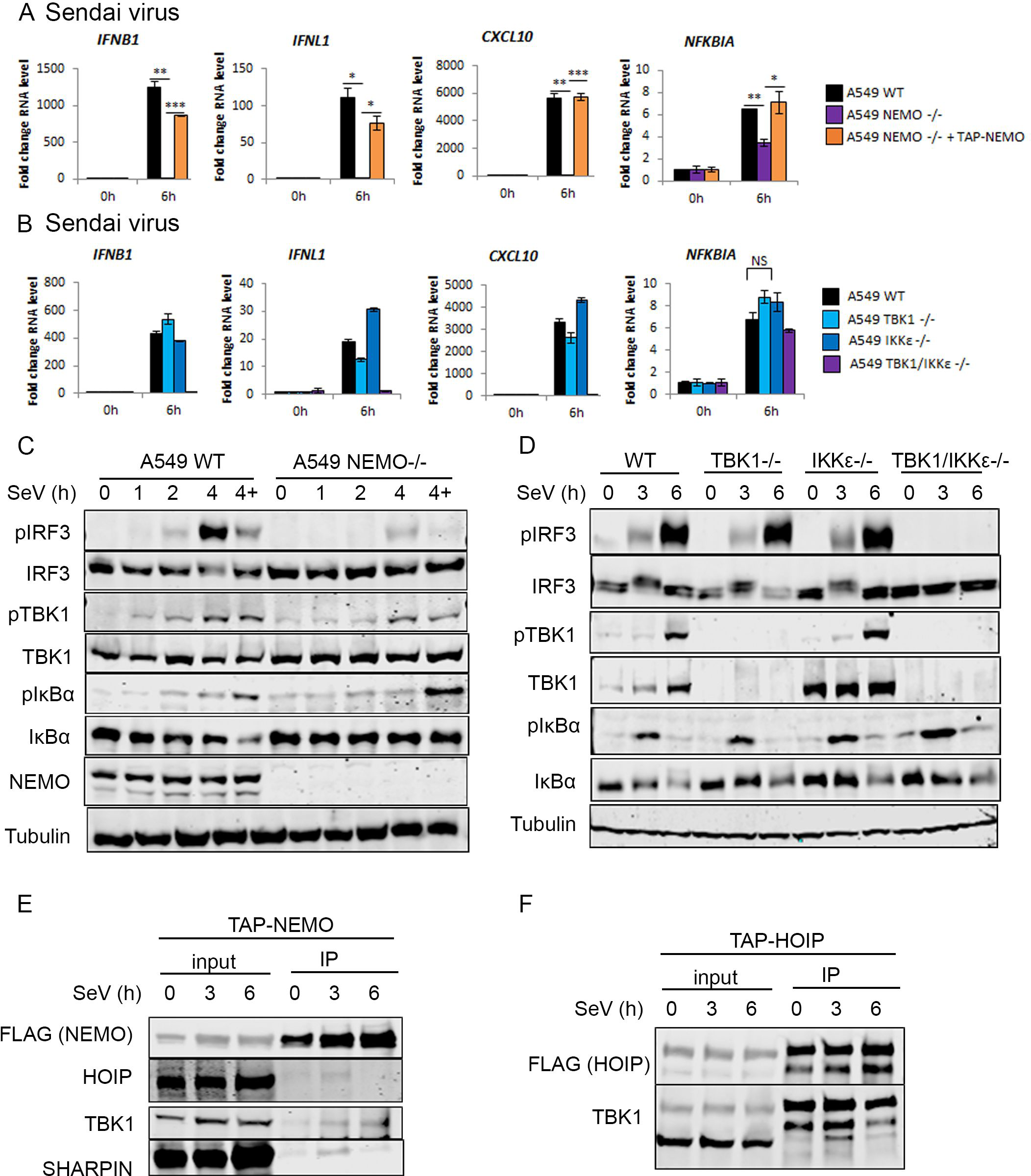
LUBAC interacts with TBK1 and NEMO downstream of RIG-I activation. qPCR to measure transcription of indicated genes during SeV infection at 1:300 dilution in A) A549 WT, NEMO -/-, NEMO -/- + TAP-NEMO cells and B) A549 WT, TBK1 -/-, IKKε -/- andTBK1/IKKε -/- cells. Western blotting analysis of signalling protein phosphorylation during SeV infection at 1:300 dilution in A) A549 WT, NEMO -/-, NEMO -/- + TAP-NEMO cells and B) A549 WT, TBK1 -/-, IKKε -/- andTBK1/IKKε -/- cells. Western blotting analysis of Flag-M2 IP in E) A549 TAP-NEMO and F) A549 TAP-HOIP-WT cells infected with SeV at a 1:300 dilution

### LUBAC interacts with TBK1 and NEMO downstream of RIG-I

Since the E3 ligase activity of HOIP is only partially required for LUBACs function in RIG-I signalling, we explored the possibility that LUBAC plays a structural role in the RIG-I signalling complex. Isolation of the endogenous RIG-I/MAVS signalling complex is complicated by low levels of RIG-I protein expression and its localisation at the mitochondria. We therefore used the TAP-NEMO and TAP-HOIP rescue cell lines to immunoprecipitate protein complexes following SeV infection. At 3 hours post SeV infection, HOIP, SHARPIN and TBK1 could all be found in complex with immunoprecipitated NEMO, enriched compared to mock-infected cells (Fig. 7E). By 6 hours post infection, only TBK1 remained in complex with NEMO (Fig. 7E). Similarly, HOIP was found to co-immunoprecipitate with TBK1 in an SeV-dependent manner (Fig. 7F). As such, LUBAC is specifically and transiently recruited to NEMO and TBK1 in an SeV-infection-dependent manner, consistent with the requirement for HOIP protein and E3 ligase activity in RIG-I signalling. Overall these results define HOIP and HOIL-1 as critical components of the RIG-I signalling complex required for anti-viral innate immunity.

## Discussion

During RNA virus infection RIG-I is activated by viral RNAs and undergoes a conformational switch allowing the construction of a large, multiprotein signalling complex, which relies on post translational modification of component proteins. Multiple E3 ubiquitin ligases and kinases regulate this dynamic process to generate optimal signalling outputs in a given cellular context. The M1 ubiquitin E3 ligase LUBAC modulates signalling outputs of multiple immune SCs, amplifying gene activation and regulating programmed cell death signalling outputs^13^. During TNFR1 signalling, LUBAC is recruited by binding to the K63-linked ubiquitin chains produced by cIAP1/2. LUBAC then adds M1-linked ubiquitin chains to RIP1, NEMO, TNFR1 and TRADD^14,30,31^, as well as to pre-established K63-linked chains, generating K63-/M1-linked heterotypic chains^32,33^. This results in the formation of the TNFR1-SC, also known as complex I of TNFR1 signalling and increased recruitment of NEMO ^34^. LUBAC functions similarly in other immune signalling pathways, conjugating linear ubiquitin chains to other targets including RIPK2, TRADD, TNFR1 itself, IRAK1/2/4 and MyD88^35^. LUBAC E3 ligase activity can also generate M1-/K63-linked heterotypic chains, conjugated to NEMO in IL-1β and TLR3 signalling, RIPK1 in TLR3 signalling, and RIPK2 in NOD2 signalling ^15,32,33,36^.

Here we define specific and separate contributions of LUBAC components and M1 chains to RIG-I signalling using a knockout approach in a system where the transcriptional response to intracellular RNAs or infection with SeV and ZIKV was entirely dependent on RIG-I signalling, as previously reported^37–39^. This approach allowed us to determine specific RIG-I signalling outputs and the relative contributions of LUBAC components to those processes. We identified HOIP and HOIL-1 as essential components of RIG-I signalling that are required for IRF3 and NF-κB activation by RIG-I, and for the IFN-I and III response to dsRNA, ZIKV and SeV. The E3 ligase activity of HOIP is partially required RIG-I signalling, while SHARPIN is dispensable for RIG-I-driven gene activation and may negatively regulate this process in human cells. The ligase-independent function of HOIP in RIG-I signalling suggests potential for a contribution of HOIP as a scaffold or for the ligase function of HOIL-1 in activating downstream signalling. We also confirmed the essential contribution of MAVS, NEMO and TBK1/IKKε to RIG-I signalling and show that LUBAC is recruited to NEMO during SeV infection, in keeping with the requirement of HOIP and HOIL-1 for signalling activation downstream of NEMO.

Other descriptions of LUBAC in RIG-I signalling have analysed RIG-I signalling outputs in human cells overexpressing LUBAC, using siRNA knockdowns of HOIP and HOIL-1, or in murine cells expressing an incomplete HOIL-1 deletion, leading to inconsistent conclusions depending on the system^7,16,17,19,20^. Incomplete reduction or partial genetic deletion of LUBAC components may not result in the same outcome as complete deletion of the protein, and overexpression of LUBAC is known to provide conflicting positive and negative signals. Our data using cells in which HOIP or HOIL-1 are fully genetically ablated clarify the role of these proteins in RIG-I driven IRF3 and NF-kB activation and are similar to what we observed in TLR3 signalling^15^. Previous studies using cells from the SHARPIN mutant *cpdm* mouse infected with vesicular stomatitis virus concluded that LUBAC does not regulate RNA virus infection, are now clarified by our data showing that SHARPIN is not required for RIG-I signalling, even when the other components are ^16^. Our data are more consistent with that of Brazee *et al*. showing a reduced IFN response in influenza A virus-infected mice lacking HOIP or HOIL-1 in the lung epithelium^19^. The differential requirement for LUBAC components has been observed in other contexts, such in thymic development^40^, and the E3 ligase-independent functions of HOIP are also observed in B cell receptor signalling^41^. It will be interesting to understand further how SHARPIN regulates NF-κB activation in the context of multiple receptor SCs but is not required for RIG-I-driven NF-κB activation.

We propose a two-step model for how LUBAC regulates RIG-I signalling, in which HOIP and HOIL-1 act as scaffolds to allow proper formation of the RIG-I signalling complex, before HOIP conjugates linear ubiquitin chains within the complex to enhance and stabilise recruitment of downstream signalling proteins. We suggest that LUBAC is recruited to the RIG-I signalling complex by binding K63-ubiquitin chains, upon which it recruits/activates TBK1 and conjugates M1 chains to NEMO or (an)other target(s), further enhancing recruitment of NEMO, LUBAC and other M1-binding proteins, thereby amplifying downstream signalling. M1-ubiquitin chains do not appear to be required for the role of LUBAC in regulating either TBK1 or IRF3 activation, or the phosphorylation and degradation of IκBα to activate NF-κB, so this is only dependent on the presence of HOIP and HOIL-1 at the RIG-I-SC and not M1-ubiquitin chain formation. M1 ubiquitin chains are, however, required to enhance recruitment of signalling proteins and boost downstream responses. We suggest that this may be caused by the formation of M1/K63-linked hybrid ubiquitin chains that function to amplify IRF3 activation in the RIG-I-SC. The mechanism by which this regulation occurs also relies on our knowledge of LUBAC at the TNFR1-SC. In TNFR1 signalling, both NEMO and LUBAC are initially recruited by binding to ubiquitin chains generated by cIAPs^14,42^. LUBAC then adds M1 ubiquitin chains to various components of the TNFR1-SC^14,30,31,43^, including TRADD and RIP1, which enhances recruitment and retention of NEMO, which a much higher affinity for M1-ubiquitin chains than K63/K11-linked chains^30,34,44^. The recruitment of TBK1 and IKKε to the TNFR1-SC is also mediated largely by M1 ubiquitin chains, as well as TANK and NAP1^31,45^. Similarly, TRAF proteins have been shown to produce K63-ubiquitin chains that recruit NEMO to the RIG-I signalling complex. Therefore, we propose that K63-ubiquitin chains generated by TRAFs recruit LUBAC and NEMO to the RIG-I signalling complex, and that the presence of both LUBAC and NEMO here enables recruitment and activation of TBK1/IKKε and IRF3, as well as NF-κB. Overall our data adds detail to the significant contribution of LUBAC to anti-viral immunity and places HOIP and HOIL-1, but not SHARPIN, as key regulators of the IFN response to infection by RNA viruses.

## Acknowledgements

This work was funded by a Wellcome Trust PhD studentship 203778/Z/16/Z to HCT and BF, a UKRI/BBSRC research project grant BB/S001336/1 to BF, Conselho Nacional de Desenvolvimento Cient fico e Tecnol gico (CNPq) grants 307889/2020-3 and 407609/2018-0 to DSM, a Cancer Research UK Programme Grant (A17341), a Wellcome Trust Investigator Award (214342/Z/18/Z), a Medical Research Council Grant (MR/S00811X/1), two collaborative research centre grants (CRC 1399, C06, and SFB1403–414786233) funded by the Deutsche Forschungsgemeinschaft (DFG) and an Alexander von Humboldt Foundation Professorship awarded to HW.

## Conflict of interest

The authors declare no conflict of interest.

## Data availability

The datasets used during the current study are available from the corresponding author on reasonable request. All data generated or analysed during this study are included in this published article and its supplementary information files.

## Author Contributions

BJF, HT, HW, NI and DM performed study concept and design; HT, BJF, HW, ER, DM, NI and DM performed development of methodology and writing, review and revision of the paper; HT, CL, ER, DM, DP, MO, and BJF provided acquisition, analysis and interpretation of data, and statistical analysis; All authors read and approved the final paper.

## Supplementary data

**Supplementary Table 1.**
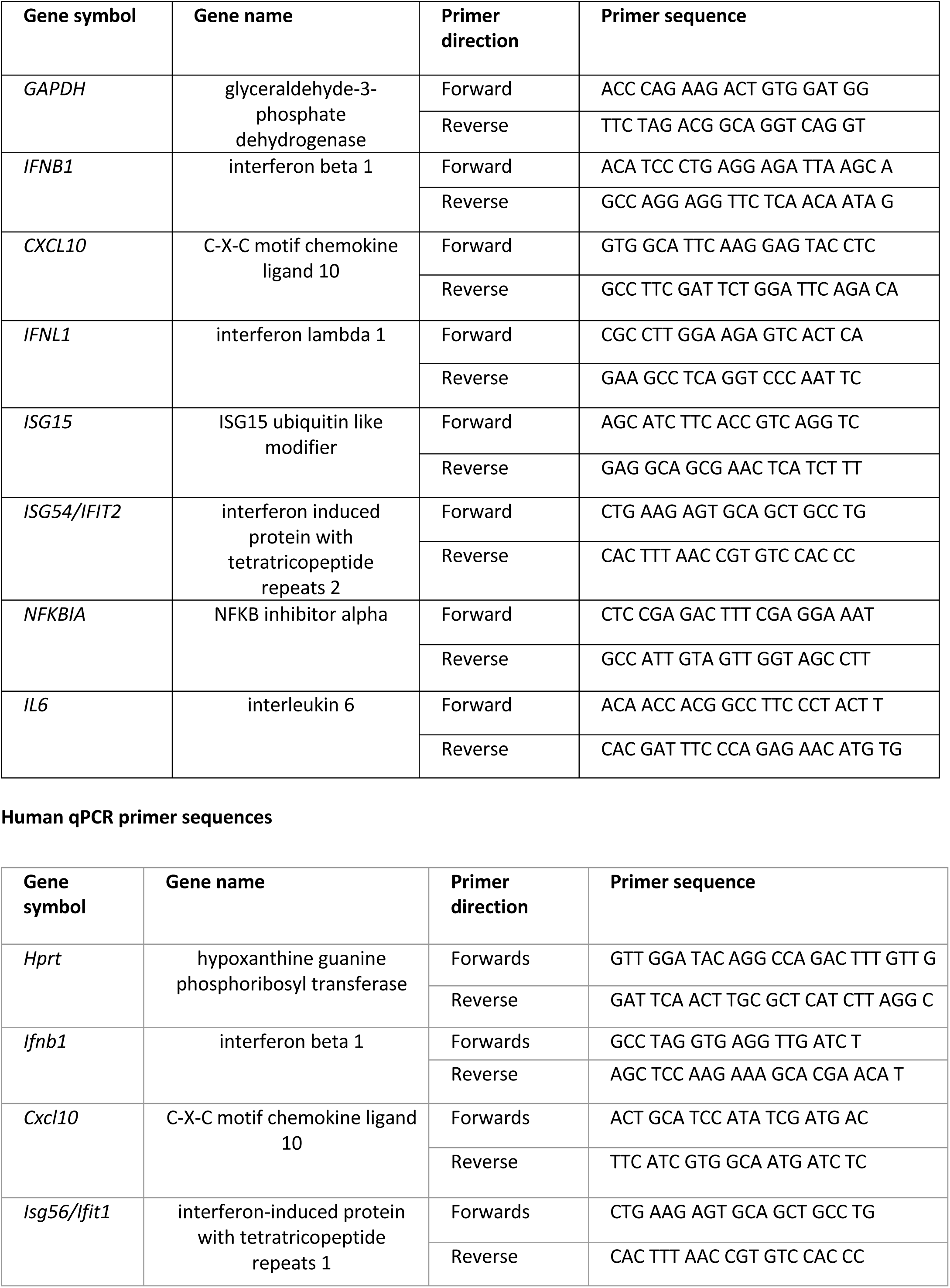
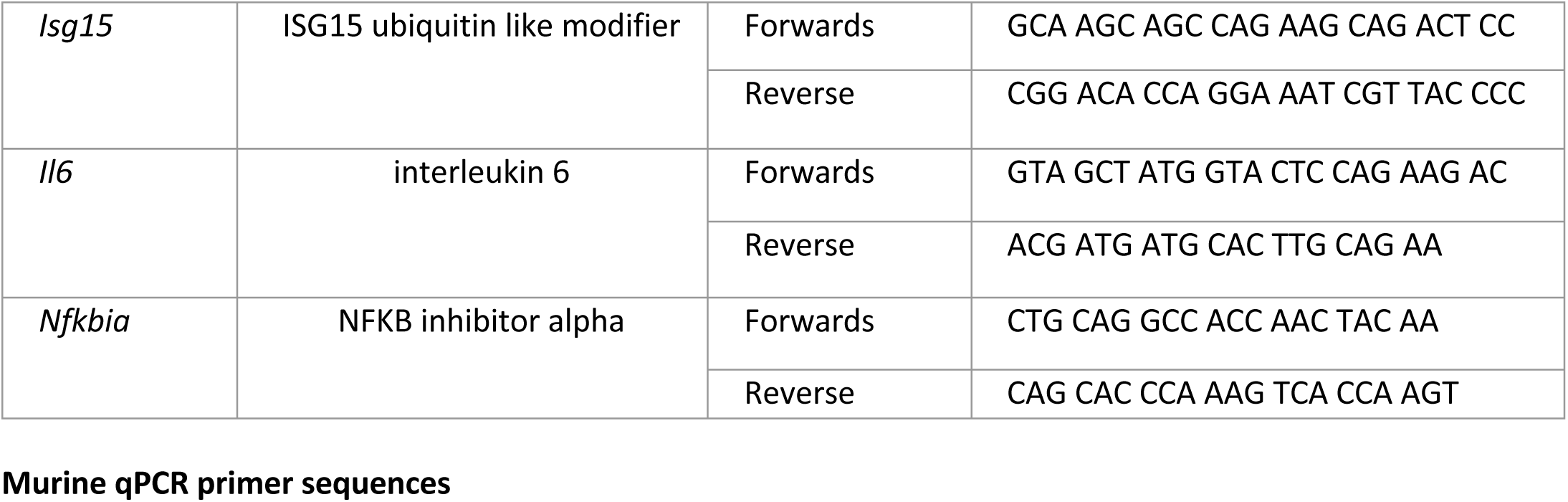
qPCR primer sequences used in this study.

**Supplementary Table 2.**
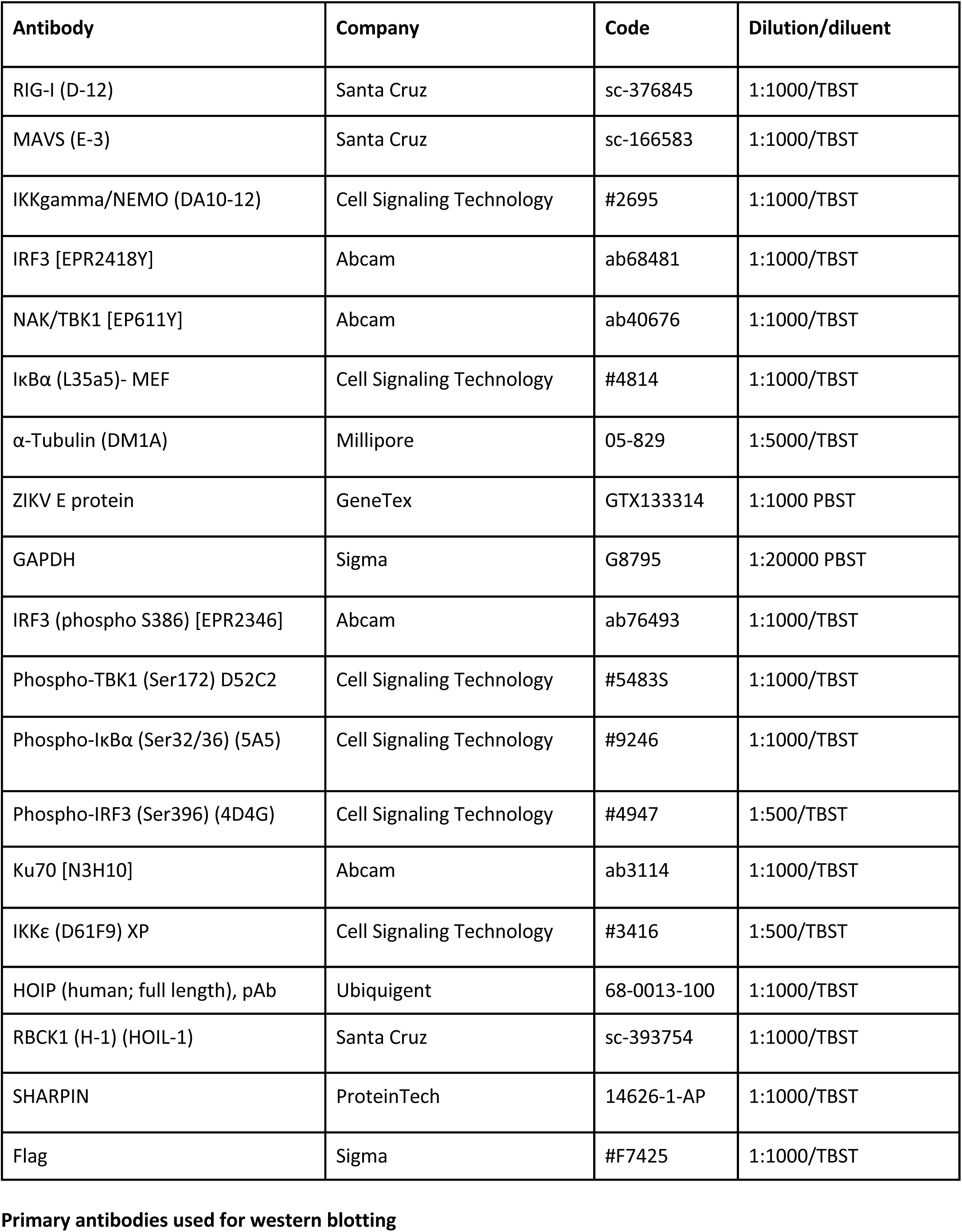
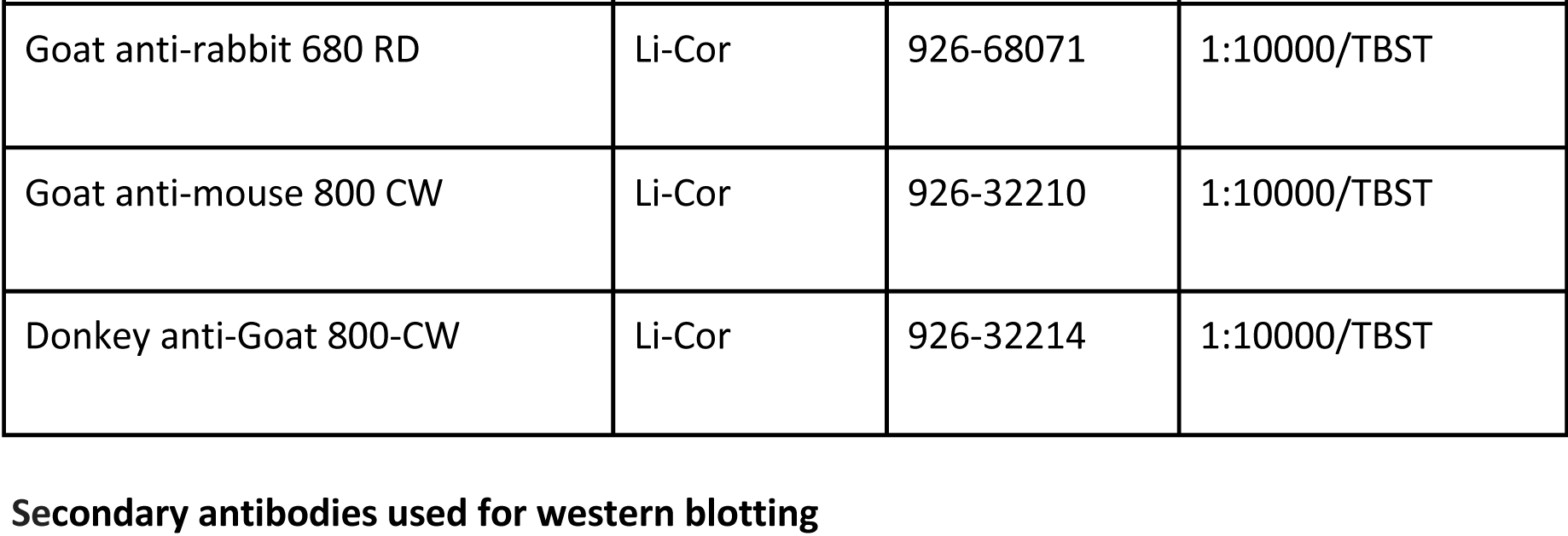
Primary and secondary antibodies used for immunoblotting in this study.

**Supplementary Table 3.**
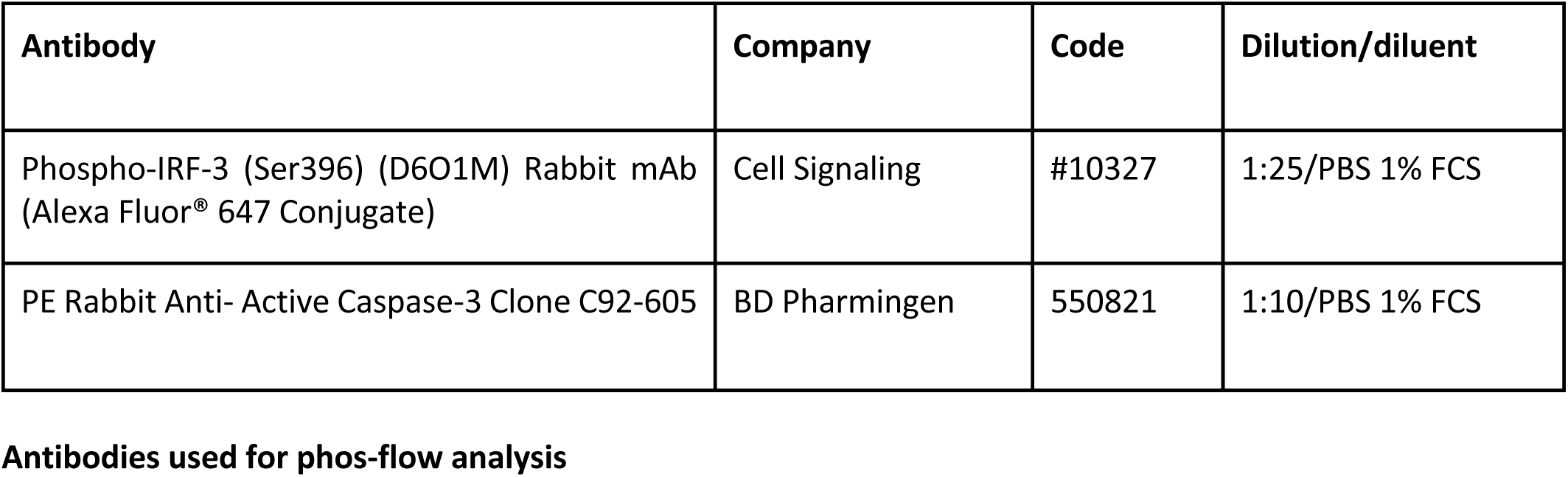
Antibodies used for PhosFlow analysis in this study.

| **Antibody** | **Company** | **Code** | **Dilution/diluent** |
| --- | --- | --- | --- |
| Phospho-IRF-3 (Ser396) (D6O1M) Rabbit mAb (Alexa Fluor® 647 Conjugate) | Cell Signaling | #10327 | 1:25/PBS 1% FCS |
| PE Rabbit Anti- Active Caspase-3 Clone C92-605 | BD Pharmingen | 550821 | 1:10/PBS 1% FCS |

**Supplementary Table 4.**
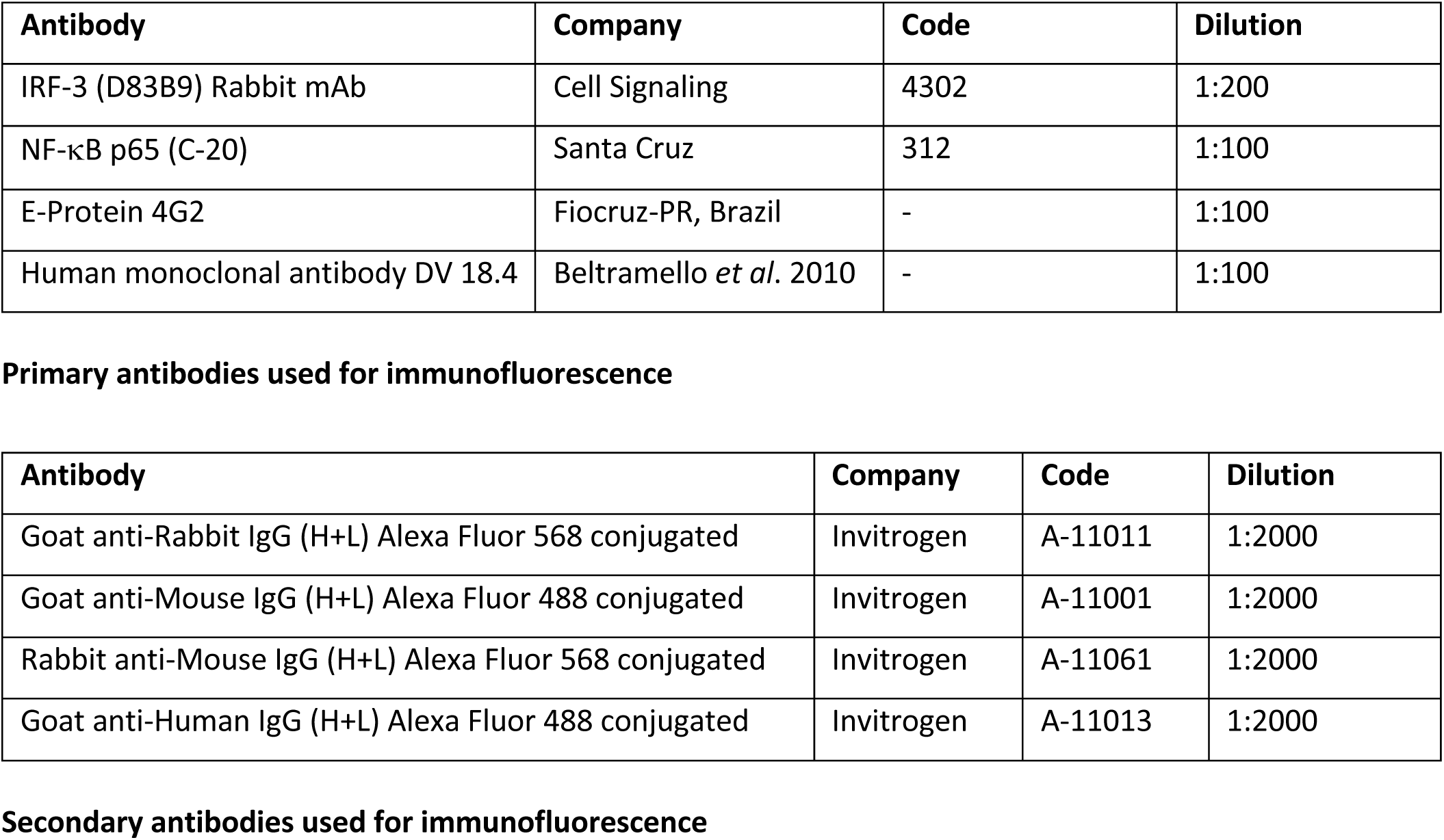
Antibodies used for immunofluorescence analysis in this study.

**Supplementary Figure S1:**
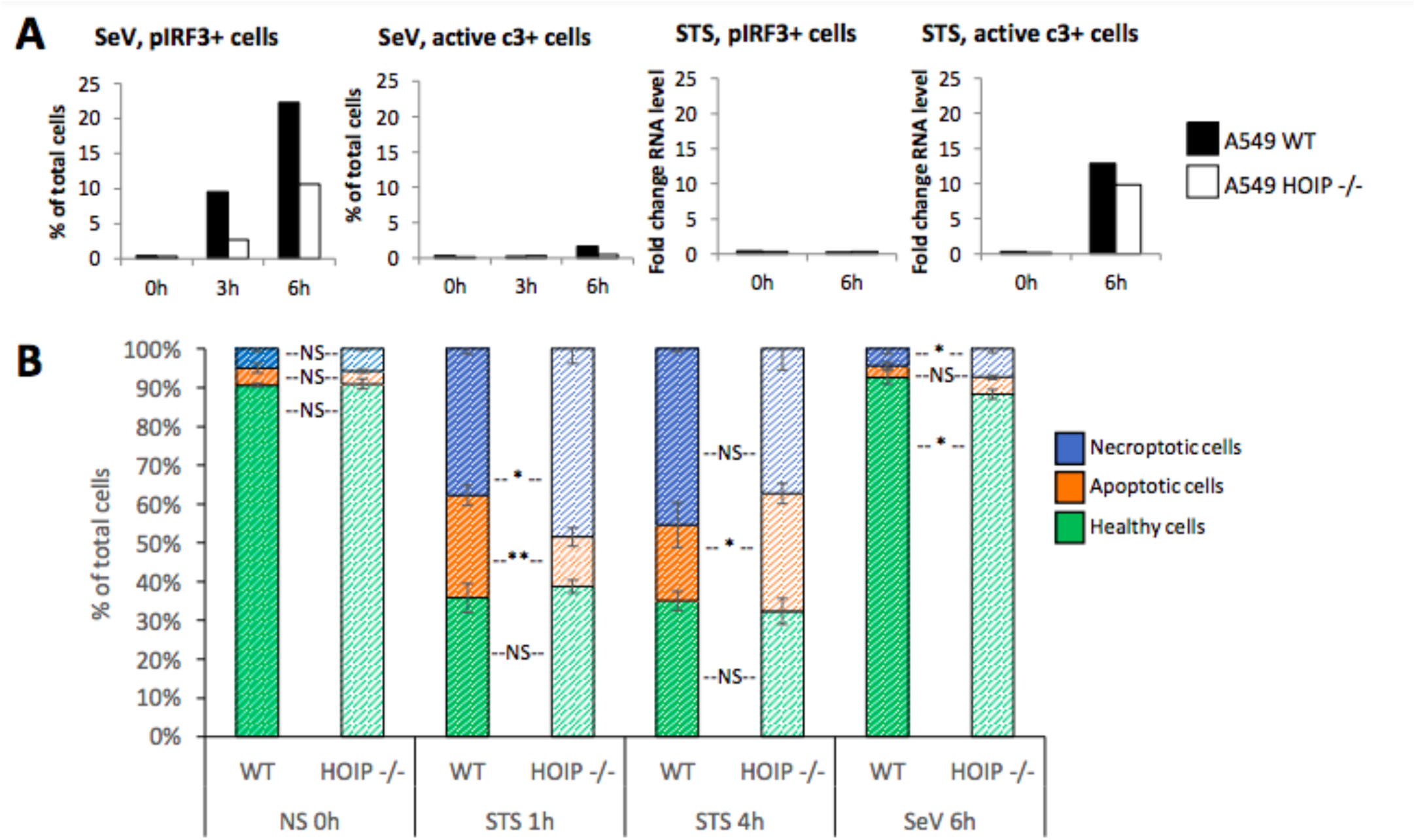
Loss of HOIP does not result in RIG-I-driven cell death in A549 cells. A) Phos-flow to measure cells expressing phospho-IRF3 and cleaved caspase 3 in A549 WT and HOIP -/- cells infected with SeVat 1:300 dilution. B) Nucleocounter (NC-250) Vitality Assay in A549 WT and HOIP -/- cells treated with staurosporine at 2 μM or infected with SeV at 1:300 dilution for the indicated times. NS = not stimulated.

**Supplementary Figure S2:**
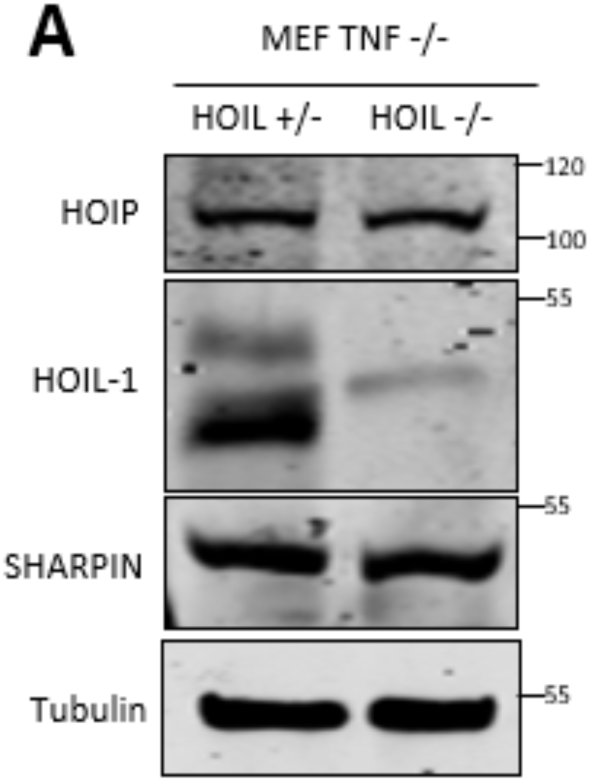
Western blotting analysis of LUBAC components in MEF TNF -/- HOIL +/- and TNF -/- HOIL -/- cells.

**Supplementary Figure S3:**
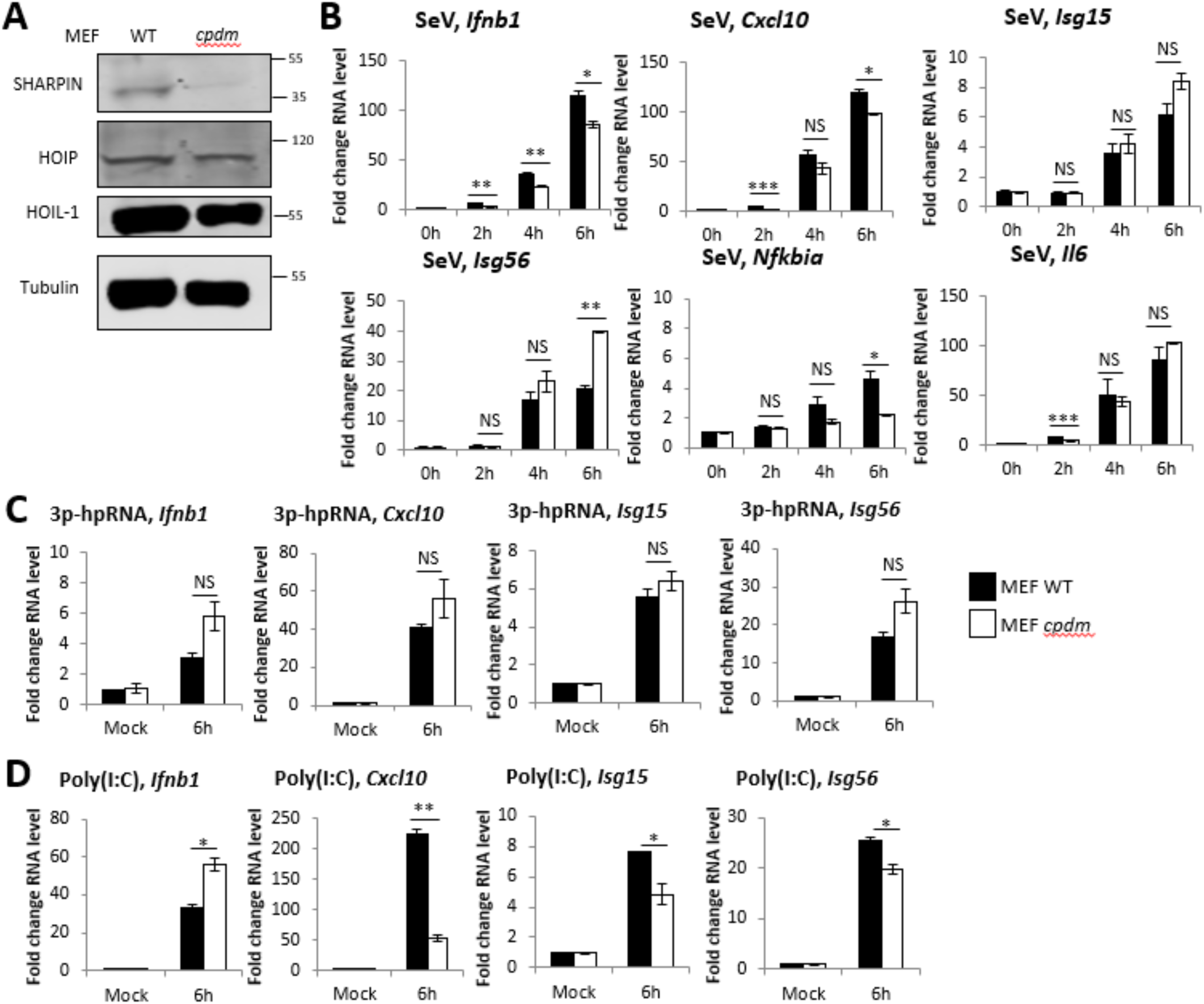
SHARPIN is not required for RIG-I immune response to SeV and synthetic RNAs in MEF cells. A) Western blotting analysis of MEF WT and *cpdm* cells. qPCR to measure transcription of indicated genes in MEF WT and *cpdm* cells B) infected with SeV at a 1:300 dilution or transfected with C) 1 μg 3p-hpRNA and D) 1 μg Poly(I:C).

**Supplementary Figure 4:**
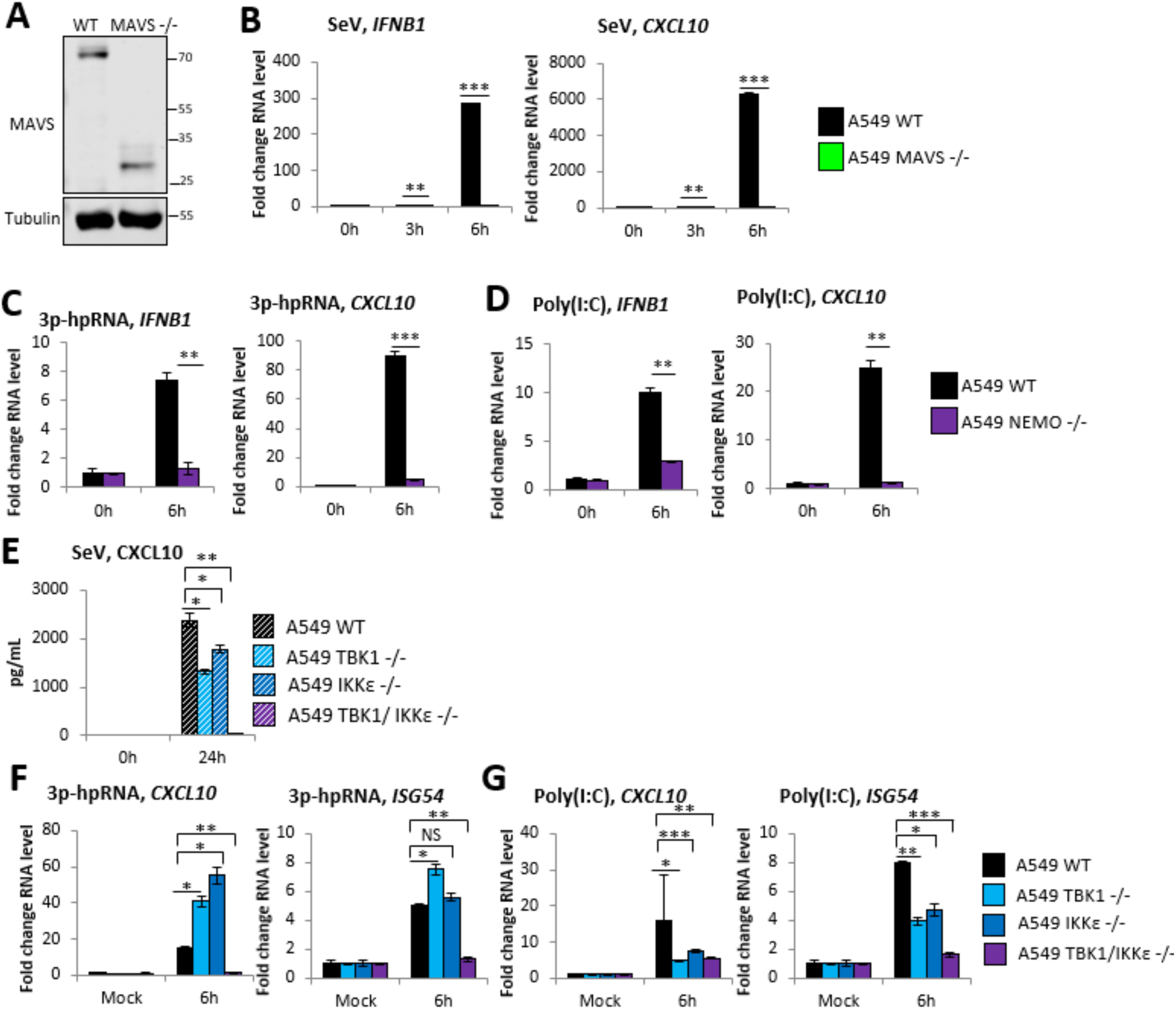
Requirement of TBK1, IKKε, NEMO and MAVS in RIG-I signaling. A) Western blotting analysis of A549 WT and MAVS -/- cells. B) Transcription of indicated genes measured by qPCR in A549 WT and MAVS -/- cells infected with SeV at 1:300 dilution. qPCR to measure transcription of indicated genes in A549 WT and NEMO -/- cells transfected with C) 1 μg 3p-hpRNA and D) 1 μg Poly(I:C). A549 WT, TBK1 -/-, IKKε -/- and TBK1/IKKε -/- cells E) infected with SeV at 1:300 and ELISA to measure CXCL10 secretion, transfected with F) 1 μg 3p-hpRNA and G) 1 μg Poly(I:C) and qPCR to measure transcription of indicated genes.

## Notes

### Competing Interest Statement

The authors have declared no competing interest.

